# Excess folic acid exposure increases uracil misincorporation into DNA in a tissue-specific manner in a mouse model of reduced methionine synthase expression

**DOI:** 10.1101/2024.06.20.599913

**Authors:** Katarina E. Heyden, Olga V. Malysheva, Amanda J. MacFarlane, Lawrence C. Brody, Martha S. Field

**Author notes:** Corresponding author: Martha S. Field, Cornell University.

## Abstract

**Background:** Folate and vitamin B12 (B12) are cofactors in folate-mediated one-carbon metabolism (FOCM), a metabolic network that supports synthesis of nucleotides (including thymidylate, or dTMP) and methionine. FOCM impairments such as a deficiency or imbalance of cofactors can perturb dTMP synthesis, causing uracil misincorporation into DNA.

**Objective:** The purpose of this study was to determine how reduced expression of the B12-dependent enzyme methionine synthase (MTR) and excess dietary folic acid interact to affect folate distribution and markers of genome stability in mouse tissues.

**Methods:** Heterozygous *Mtr* knockout mice (*Mtr*^*+/-*^) model the FOCM-specific effects of B12 deficiency. Folate accumulation and vitamer distribution, genomic uracil levels, and phosphorylated histone γH2AX immunostaining were measured in male *Mtr*^*+/+*^ and *Mtr*^*+/−*^ mice weaned to either a folate-sufficient control (C) diet (2 mg/kg folic acid) or a high folic acid (HFA) diet (20 mg/kg folic acid) for 7 weeks.

**Results:** Exposure to the HFA diet led to tissue-specific patterns of folate accumulation, with plasma, colon, kidney, and skeletal muscle exhibiting increased folate concentrations compared to control. Liver total folate did not differ. Though unmetabolized folic acid (UMFA) increased 10-fold in mouse plasma with HFA diet, UMFA accounted for less than 0.2% of total folate in liver and colon tissue. Exposure to HFA diet resulted in a shift in folate distribution in colon tissue with higher 5-methyl-THF and lower formyl-THF than in control mice. *Mtr* heterozygosity did not impact folate accumulation or distribution in any tissue. Mice on HFA diet exhibited higher uracil in genomic DNA and γH2AX foci in colon. Similar differences were not seen in liver.

**Conclusions:** This study demonstrates that folic acid, even when consumed at high doses, does not meaningfully accumulate in mouse tissues, although high-dose folic acid shifts folate distribution and increases uracil accumulation in genomic DNA in colon tissue.

## Introduction

The relationship between folate and vitamin B12 (B12) in supporting nucleotide synthesis has been appreciated for decades (1). Folate and B12 are required cofactors in folate-mediated one-carbon metabolism (FOCM), a metabolic pathway that provides one-carbon donors for nucleotide biosynthesis and methyl-donor production. In mammals, only two enzymes in the body require B12: methylmalonyl CoA mutase (MCM), a mitochondrial enzyme which aids in amino acid and fatty acid metabolism via the tricarboxylic acid cycle, and methionine synthase (MTR), a cytosolic enzyme which is part of FOCM (2). MTR requires both folate and B12 as cofactors to catalyze the regeneration of methionine from homocysteine. In this two-step process, 5-methyl-tetrahydrofolate (5-methyl-THF) provides a methyl group to cobalamin (a form of B12) to produce methylcobalamin and releases THF (3). Methylcobalamin then donates its methyl group to homocysteine to regenerate methionine. Conversion of 5,10-methylene-THF to 5-methyl-THF by methylenetetrahydrofolate reductase (MTHFR) is irreversible and MTR is the only enzyme capable of metabolizing 5-methyl-THF to release THF (4) (**Fig 1**). Release of THF is crucial, as it is the bioactive form of folate required to carry the one-carbon groups for nucleotide synthesis. When MTR activity is compromised as happens in B12 deficiency, cellular folate accumulates as 5-methyl-THF during, creating the hallmark “5-methyl-THF” trap. This reduces the availability of other folate cofactors to support nucleotide synthesis. Eventually megaloblastic anemia presents because dimished nucleotide pools are unable to support DNA synthesis and cell division. Slowed red blood cell production then results in a macrocytic anemia.

**Figure 1.**
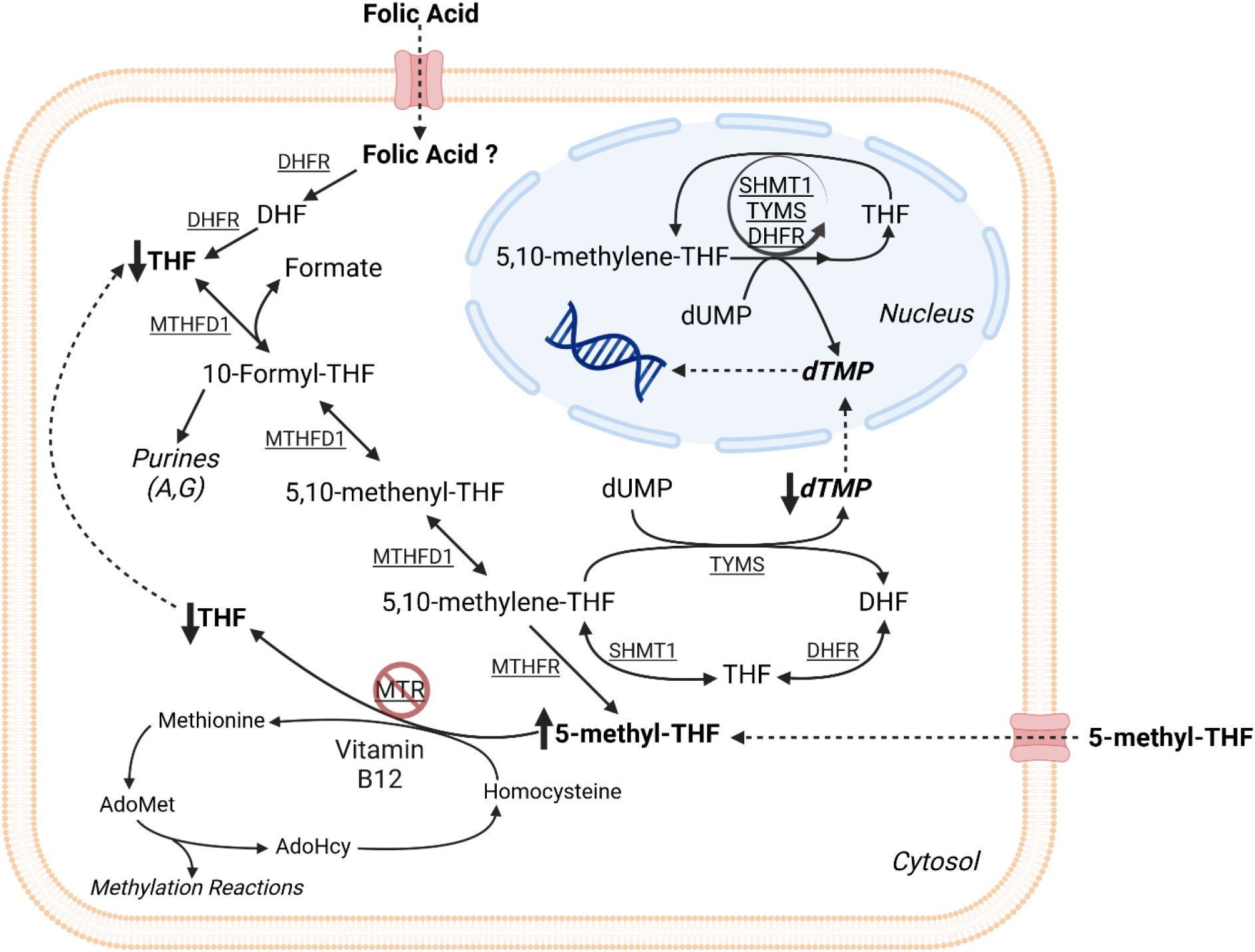
The effects of reduced *Mtr* expression and excess folic acid on cellular folate-mediated one carbon metabolism (FOCM) Partial knockout of the *Mtr* gene in mice models vitamin B12 deficiency specific to the FOCM pathway. Reduced Mtr expression causes elevation of cellular folate as 5-methyl-THF and corresponding decrease of folate as THF. The conversion of 5,10-methylene-THF to 5-methyl-THF is irreversible and 5-methyl-THF can only be metabolized to THF by the MTR enzyme. AdoHcy, S-adenosylhomocysteine; AdoMet, S-adenosylmethionine; DHFR, dihydrofolate reductase; dTMP, deoxythymidine monophosphate; dUMP, deoxyuridine monophosphate; MTHFD1, methylenetetrahydrofolate dehydrogenase 1; MTHFR, methylenetetrahydrofolate reductase; MTR, methionine synthase; SHMT, serine hydroxymethyltransferase; THF, tetrahydrofolate; TYMS, thymidylate synthase.

It is important to note that folic acid is chemically distinct from naturally occurring folate. While folic acid appears to fill the same biological role as natural folate, it is unknown as to whether it is associated with other biochemical changes. Folic acid is a synthetic, oxidized folate form commonly used in grain fortification and dietary supplements, while natural folates forms are reduced. There is no Tolerable Upper Intake Level (UL) for reduced folates found in food (THF forms, Figure 1), but there is a UL for folic acid consumption. Folic acid does not readily participate in FOCM but must first be metabolized into a bioactive form of folate. Dihydrofolate reductase (DHFR), the enzyme responsible for converting DHF to THF (Figure 1), is required to convert folic acid to DHF. Unmetabolized folic acid (UMFA) has been found in nearly all serum samples from American children and adults (5). This demonstrates that folic acid is not entirely metabolized upon absorption, and is commonly consumed in amounts surpassing the body’s threshold to readily metabolize it. Dietary supplements are the main contributors to high folate status and the presence of UMFA (6).

Recently, the folate-B12 interrelationship has received increased attention due to human observational data associating the combination of elevated folate status with low or deficient B12 status (known as “high folate/low B12”) with an array of negative health outcomes: worsened clinical markers of B12 deficiency, diminished response to B12 treatment, and increased risk for cognitive impairment and decline in older adults (7,8). The observation of negative health outcomes are difficult to separate from selection bias and other confounders, and such studies have not investigated biological mechanisms (9). There is little data from B12-deficient model systems to investigate the mechanistic basis for these associations.

This study used mice missing one functional copy of the gene for the B12-dependent MTR enzyme. These mice should only have 50% of MTR enzyme activity. Complete knockout of this enzyme produces early embryonic lethality (10). Mice with reduced MTR levels were exposed to excess dietary folic acid. High folic acid consumption caused tissue-specific differences in folate accumulation. In colon, high dietary folic acid perturbed cellular folate distribution towards more 5-methyl-THF, caused uracil accumulation in DNA, and modestly elevated DNA damage. Liver was unaffected. In this model, *Mtr* genotype had no effect on outcomes measured, suggesting effects from HFA are not exacerbated when combined with B12 deficiency as modeled by decreased flux through MTR.

## Methods

### Animal breeding and dietary intervention

All mice were maintained under specific-pathogen-free conditions in accordance with standard of use protocols and animal welfare regulations. All study protocols were approved by the Institutional Animal Care and Use Committee of Cornell University. *Mtr*^*+/-*^ male mice, generated as previously described (10), were backcrossed to C57Bl/6J for over 10 generations and crossed to C57BL/6J female mice. Male *Mtr*^*+/+*^ and *Mtr*^*+/-*^ offspring were weaned and randomized to be fed one of two defined diets at three weeks of age. The diets consisted of defined AIN93G (11) control (C) diet containing 2 mg/kg folic acid (#117814GI; Dyets, Inc., Bethlehem, PA), or an AIN93G-based high folic acid (HFA) diet containing 20mg/kg folic acid (#117876GI; Dyets, Inc., Bethlehem, PA). Dietary intake and body weight of the mice were recorded every 14 days. Mice were maintained on the assigned diets for 7 weeks and sacrificed by cervical dislocation following CO_2_ euthanasia. A subset of mice were fasted for 12 hours prior to harvest (fasted indicated in figure legend where applicable) to determine the effects of fasting on plasma and tissue folate accumulation. Whole blood was collected via cardiac puncture into Heparin-coated tubes. Plasma and red blood cells were separated by centrifugation at 2500 x g and immediately flash frozen in liquid nitrogen. Tissues were harvested, rinsed in ice-cold 1X phosphate buffered saline (PBS, Corning), and immediately flash frozen in liquid nitrogen or preserved in O.C.T. Compound (Tissue-Tek, #4583) for cryo-sectioning. All samples were then stored at -80°C for further analysis.

### *Lactobacillus casei* assay to quantify total folate concentrations

Total folate concentrations in plasma and tissues were quantified by *Lactobacillus casei* microbiological assay as previously described (12), with bacterial growth quantified at 600 nm on Epoch Microplate Spectrophotometer (Biotek Instruments). Tissue folate concentrations were normalized to protein concentration, as assessed by Lowry-Bensadoun assay (13).

### Folate distribution

Folate concentrations and vitamer form distribution were quantified by LC-MS electrospray tandem MS adapted from previously described methods (14–17). Liver and colon folate concentrations were normalized to sample weight. Plasma samples were pooled (4 mice per sample, n=3-4 samples per group) before analysis.

### Immunoblotting

Tissues were suspended in lysis buffer (150 mM NaCl, 10 mM Tris-Cl, 5 mM EDTA pH 8, 5 mM dithiothreitol, 1% Triton X-100, protease inhibitor), sonicated, and centrifuged at 4°C for 10 minutes at 14,000 x g to remove the insoluble fraction. Protein concentrations of tissue lysate supernatants were assessed by Pierce BCA Protein Assay Kit (Thermo Scientific). Samples were boiled with SDS-PAGE sample loading buffer (6X SDS) and 20 ug of protein per well was loaded on a 12% Tris-glycine SDS-PAGE gel. Proteins were transferred to a PVDF membrane (Millipore). Membranes were blocked with 5% (wt/vol) non-fat dry milk in 1X phosphate buffered saline with 0.1% Tween-20 (PBST) for 1 hour. Membranes were then incubated overnight at 4°C with primary antibody (GAPDH, 1:50,000 dilution, Cell Signaling #2118; α-MTR, 1:1,000 dilution, ProteinTech #25896; DHFR, 1:500 dilution, Cell Signaling #45710) in 5% bovine serum albumin (BSA) 0.02% NaN_3_ in PBST. Membranes were washed 3 times in PBST and incubated for 1 hour in HRP-conjugated secondary antibody diluted 1:20,000 in 5% (wt/vol) non-fat dry milk in PBST. Membranes were exposed to chemiluminescent substrate (BioRad) and visualized by ProteinSimple Imager. ImageJ software was used to quantify band intensity and protein expression.

### Determination of genomic uracil content

DNA was isolated using High Pure PCR Template Preparation Kit (Roche) per manufacturer’s protocol. Following RNAse A treatment, DNA was purified again using High Pure PCR Purification Kit (Roche) per manufacturer’s protocol, and DNA concentrations were quantified by Qubit (Thermo Scientific). 2 µg of DNA was treated with uracil DNA glycosylase (UDG, New England Biolabs, Inc.) for 60 minutes at 37°C with gentle shaking. Samples were derivatized and uracil levels quantified by gas chromatography-mass spectrometry as previously described (18).

### Immunohistochemistry

Excised tissues were stored in OCT and sectioned using a Leica cryostat CM1950. Liver (15µM) and colon (18µM) sections were placed on microscope slides and stored at -20°C prior to fixiation and staining. Tissues were fixed (10 minutes) in 4% (vol/vol) paraformaldehyde, washed (4X) with PBS, incubated (37°C for 10 min) in PBS with 0.5% Triton. Anti-γH2AX antibody conjugated to Alexa Fluor 555 (Millipore #05-636-AF555) diluted 1:1,000 in PBS with 0.5% Triton was added and slides incubated overnight (4°C). Post-binding slides were washed 4 times with PBS. DRAQ5 DNA stain (Thermo Scientific #62251) was diluted 1:1,000 in PBS, added to slides for 5 min at room temperature. Coverslips (#1.5) were mounted onto microscopy slides with 1-2 drops of Fluoromount G (Southern Biotech). All slides for each experiment were prepared simultaneously and microscopy and quantification were performed by an investigator blinded to diet and genotype. Representative images (8-bit, n=3 per slide, 3 slides per animal, n=3-5 animals per group) were acquired in one imaging session using a Zeiss LSM 710 Confocal Microscope. To allow for quantitation of γH2AX signal intensity, colon images were captured using laser power 2% and gain 610 and liver images were captured using laser power 2% and gain 640. Fiji (ImageJ) imaging software was used for image quantification. Nuclear mask creation and counts for the draq5 channel were generated as follows: Process > Smooth, Image > Adjust > Threshold (Default Algorithm), Edit > Selection > Create Selection, save nuclear area selection as a region of interest (ROI). Individual nuclei were counted manually with the multi-point tool. The nuclear ROI was then overlayed onto the 8-bit γH2Ax channel. On average, each colon image contained ∼100 nuclei and each liver image contained ∼50 nuclei. γH2AX foci counts and intensity were generated as follows: Process > Subtract Background (10 pixels, rolling paraboloid), perform difference of Gaussians using Process > Filters > Gaussian Blur, Image > Adjust > Threshold (Triangle Algorithm for colon, Minimum Algorithm for liver), Analyze > Analyze Particles (colon: size >0.1 micron^2^, liver: >0.15 micron^2^) above threshold and within nuclear ROI. Foci number, percent area, mean, and intden intensity measurements were recorded.

### Statistical analyses

All statistical analyses were performed with GraphPad Prism (version 10.1.2). Experiments comparing two genotypes (*Mtr*^*+/+*, *+/-*^) and two different diets (C/HFA) were analyzed using a two-way ANOVA with fixed effects of genotype, diet, and genotype-diet interaction, followed by Tukey’s post-hoc analysis. Body weight and food intake datasets were analyzed using a two-way ANOVA with mixed effects of diet and time. γH2AX datasets were analyzed using a two-way ANOVA with mixed effects of diet, genotype, and slide. Model assumptions of normality and equal variance for each dataset were confirmed by QQ plot and homoscedasticity plot, respectively. Data are presented as means ± standard deviation.

## Results

### HFA diet led to tissue-specific differences in folate accumulation and MTR and DHFR protein expression

Mice fed the C diet and mice fed the HFA diet did not exhibit differences in food intake or body weight over time (Fig. S1A and B). Plasma total folate concentrations were 6-fold higher in mice consuming the HFA diet (p<0.0001, Fig. 2A). The increased plasma folate concentrations were not observed in a subset of mice fasted for 12 hours (Fig. S2A). Both decreased *Mtr* expression (p<0.001) and exposure to the HFA diet (p<0.0001) resulted in lower 5-methyl-THF levels in plasma (Fig.2B). Plasma UMFA concentrations were 10-fold higher in mice consuming the HFA diet compared to those consuming the C diet (p<0.0001, Fig.2C). Total plasma folate and plasma folic acid accumulation were unaffected by *Mtr* genotype (Fig.2A, 2C), in agreement with previous findings(19).

**Figure 2.**
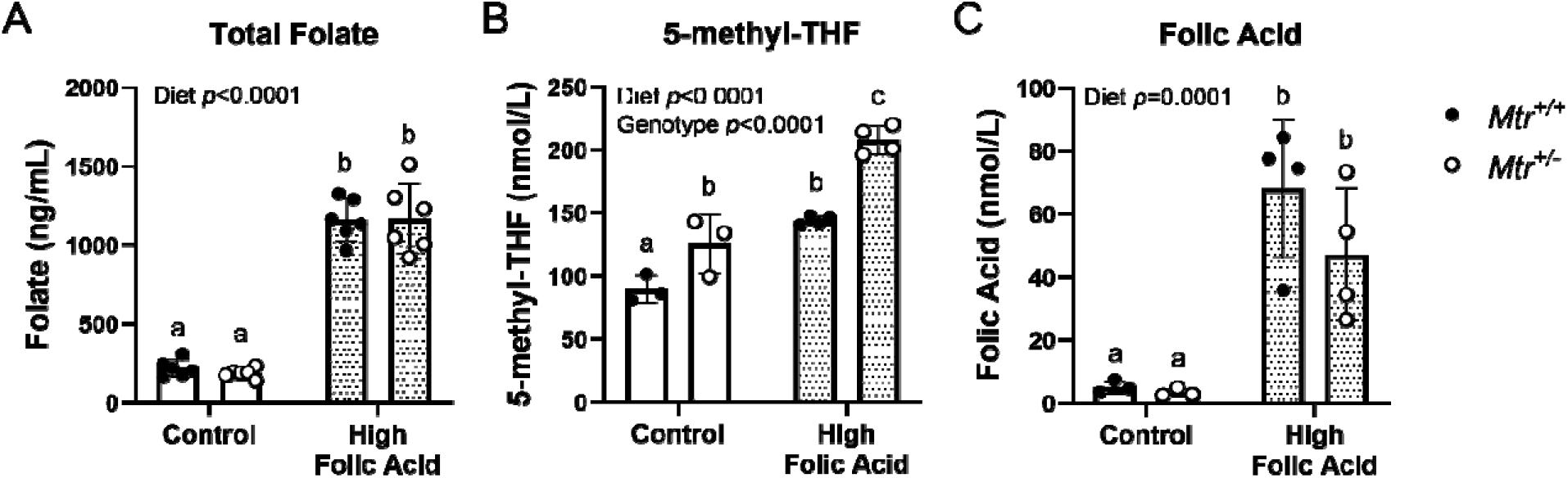
Total folate and folate form abundance in plasma of *Mtr*^*+/+*^ and *Mtr*^*+/-*^ mice fed a control or HFA diet. (A) Total folate was quantified by L. *casei* microbiological assay; n=4-6 samples per group. (B) 5-methyl-THF and (C) folic acid concentrations quantified by LCMS/MS; n=3-4 replicates per group, each replicate was pooled from 4 independent mice in that group. Two-way ANOVA with Tukey’s post hoc analysis was used to assess main effects of diet and genotype and diet–genotype interaction. Data are presented as mean ± SD with statistical significance defined *p* ≤ 0.05. Groups not connected by a common letter are significantly different. THF, tetrahydrofolate.

Exposure to the HFA diet caused changes in total folate accumulation in a tissue-specific manner (Fig 3). Liver total folate was unaffected by exposure to the HFA diet or *Mtr* genotype (Fig 3A). HFA diet exposure kidney and colon total folate was 2-fold higher in HFA fed mice (p<0.0001, Fig 3B-C), and skeletal muscle total folate was 10% higher (p<0.05, Fig 3D). Interestingly, brain total folate was 10% lower with HFA diet exposure (p<0.05, Fig 3E). Tissue total folate differences were largely consistent between nonfasted (Fig 3) and fasted mice (Supplementary Figure 2B-F). Reduction in *Mtr* expression did not impact folate accumulation in any tissue, nonfasted or fasted (Fig 3, Supplementary Fig 2).

**Figure 3.**
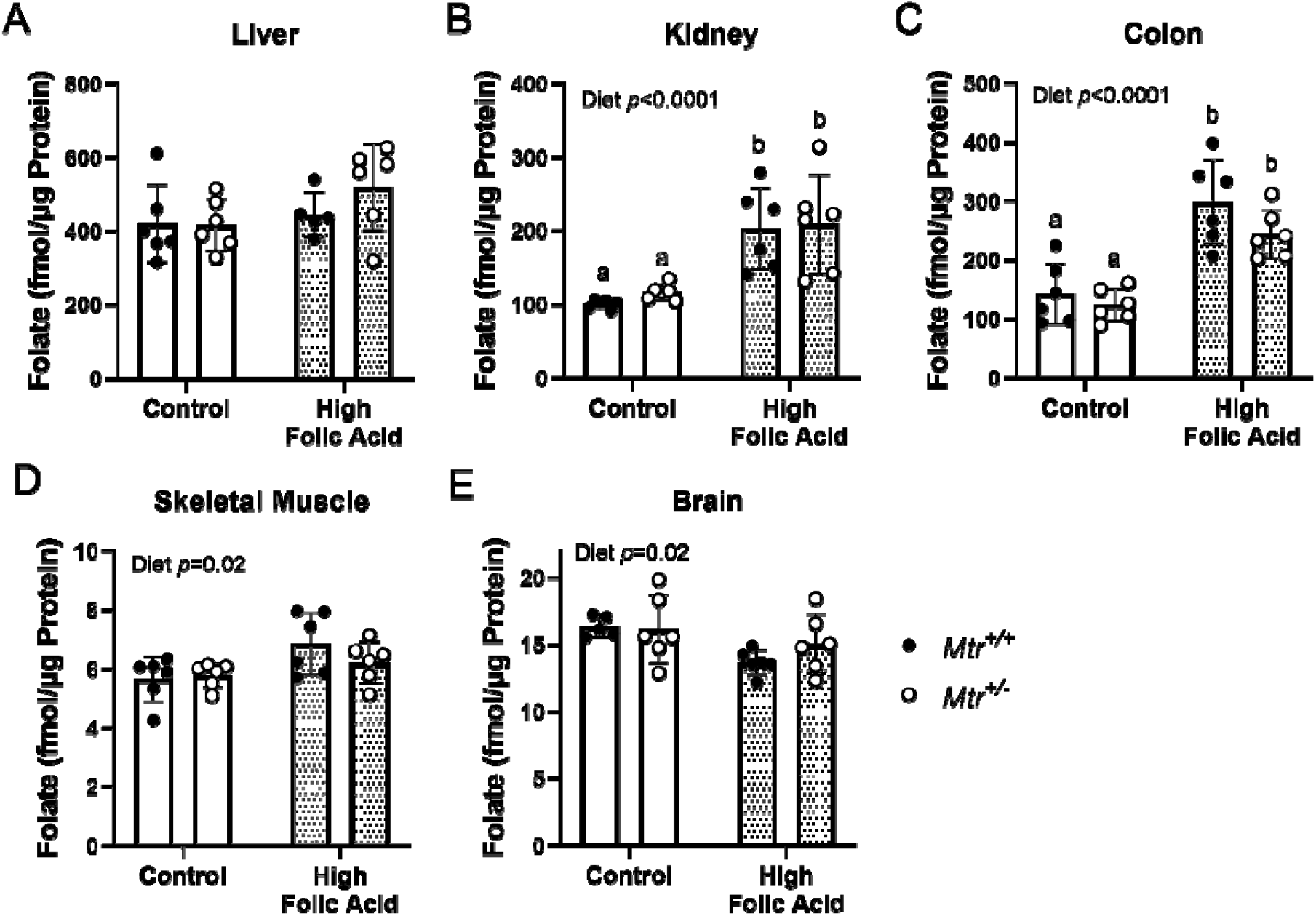
Tissue total folate accumulation in *Mtr*^*+/+*^ and *Mtr*^*+/-*^ mice fed a control or HFA diet. Total folate concentrations in (A) liver, (B) kidney, (C) colon, (D) skeletal muscle, and (E) brain were quantified by L. *casei* microbiological assay. n=6 replicates per group. Two-way ANOVA with Tukey’s post hoc analysis was used to assess main effects of diet and genotype and diet–genotype interaction. Data are presented as mean ± SD with statistical significance defined *p* ≤ 0.05. Groups that do not share a common letter are significantly different.

Liver MTR protein levels were reduced by ∼60% in *Mtr*^*+/-*^ mice (*p*<0.0001, Fig 4A-B), consistent with previous findings(19). MTR protein levels in colon, kidney, and brain were also reduced by about 50% in *Mtr*^*+/-*^ mice (*p*=0.001, *p*<0.0001, and *p*=0.004, respectively; Fig 4A-B, Supplementary Fig 3). Liver, kidney, and colon MTR protein levels were not affected by exposure to the HFA diet (Fig 4A-B). However, MTR protein levels in brain of mice consuming the HFA diet were elevated ∼2-fold compared to mice on C diet (*p*=0.02, Supplementary Fig 3D-E).

**Figure 4.**
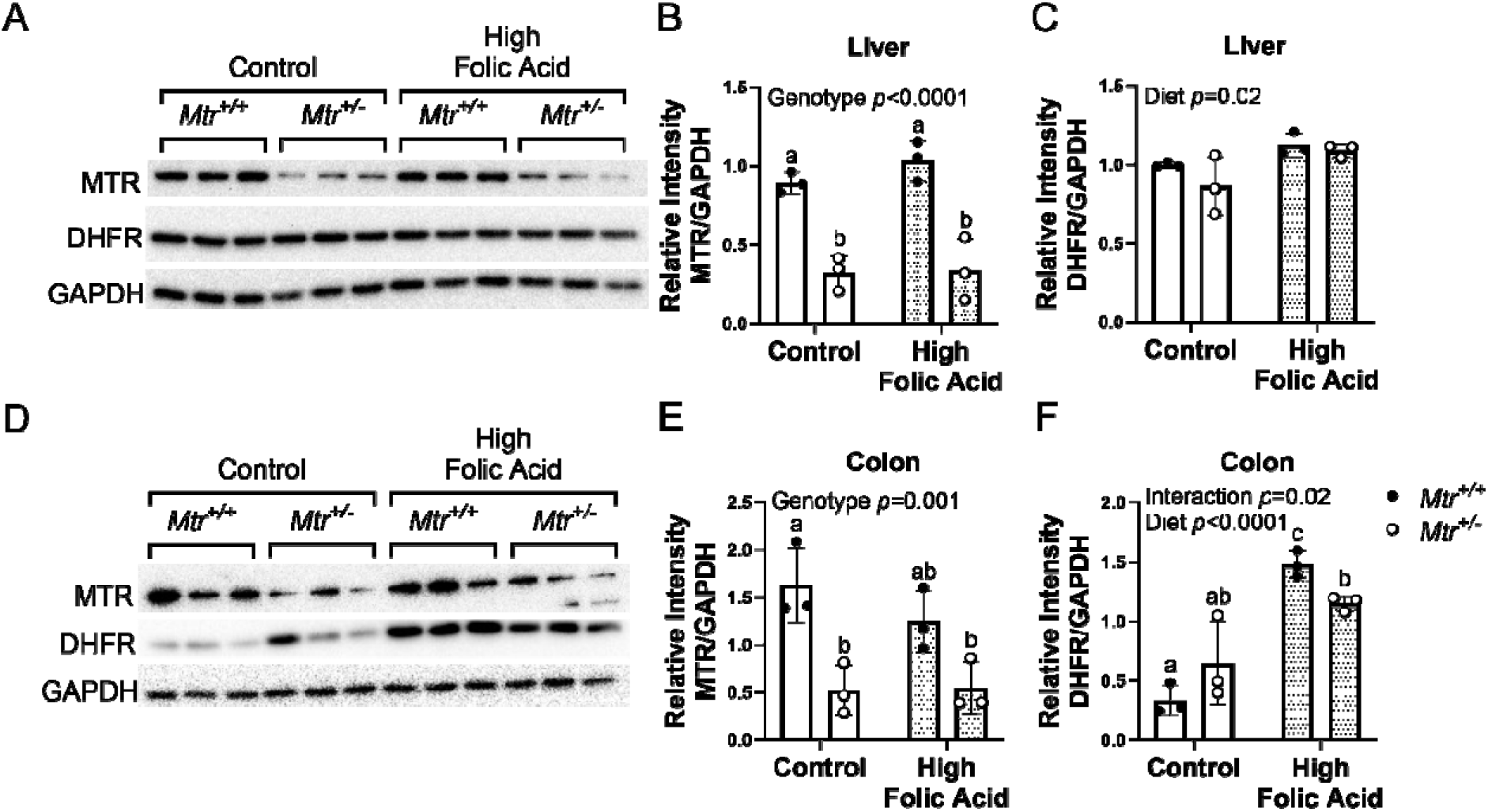
MTR and DHFR protein expression in liver and colon of *Mtr*^*+/+*^ and *Mtr*^*+/-*^ mice fed a control or HFA diet. (A) Liver Blots. (B) Liver MTR protein levels normalized to GAPDH. (C) Liver DHFR protein levels normalized to GAPDH. (D) Colon Blots. (E) Colon MTR protein levels normalized to GAPDH. (F) Colon DHFR protein levels normalized to GAPDH. n=3 per group. Two-way ANOVA with Tukey’s post hoc analysis was used to assess main effects of diet and genotype and diet–genotype interaction. Data are presented as mean ± SD with statistical significance defined *p* ≤ 0.05. Groups not connected by a common letter are significantly different. DHFR, dihydrofolate reductase; GAPDH, glyceraldehyde-3-phosphate dehydrogenase; MTR, methionine synthase.

Dihydrofolate reductase (DHFR) is the enzyme required to convert folic acid to DHF, and subsequently THF (Figure 1). To test if DHFR protein expression is altered in response to elevated folic acid consumption, protein DHFR levels were quantified by western blot. Liver and kidney DHFR protein levels did not differ by HFA diet exposure (Figure 4C and Supplementary Fig 3C). Colon DHFR protein expression was >2-fold higher in mice fed the HFA diet (*p*<0.0001, Fig 4F). A similar effect was also observed in the brain, where brain DHFR protein levels were increased by ∼50% in HFA fed mice (*p*=0.01, Supplementary Fig 3F). Reduced *Mtr* expression did not impact DHFR protein expression in liver, colon, kidney, or brain tissue.

### HFA diet altered colon folate distribution but did not increase tissue accumulation of folic acid

Previously, we have shown that folic acid does not accumulate appreciably in liver tissue of *Mtr*^*+/+*^ or *Mtr*^*+/-*^ mice consuming a defined diet with 2 mg/kg folic acid (19). To test whether a 10-fold elevation in dietary folic acid exposure would increase tissue folic acid accumulation, folate distribution was quantified in liver and colon tissue.

As shown previously (19), the most abundant forms of folate in liver were THF and 5-methyl-THF, followed by formyl-THF which made up ∼5% total folate (Fig 5, Supplementary Fig 4A-C). In agreement with *L. casei* data for total liver folate (Fig 3A), we did not observe any changes in total liver folate when measured by LCMS (Fig 5). Folic acid was detectable in liver tissue but accounted for <0.05% of all folates (Fig 5, Supplementary Figure 4D), consistent with previous observations (19,20). *Mtr*^*+/-*^ genotype resulted in elevated concentration of 5-methyl-THF in liver (*p*=0.018, Fig 5), also consistent with previous findings(19). Neither HFA diet nor *Mtr* genotype altered overall liver folate distribution (Fig 5, Supplementary Fig 4). There were minor shifts in distribution of formyl-THF and folic acid with diet and genotype (Fig 5, Supplementary Fig 4C-D). *Mtr*^*+/-*^ genotype resulted in lower formyl-THF as a proportion of total liver folate (*p*=0.01, Supplementary Fig 4C). HFA diet fed mice had higher folic acid as a proportion of total liver folate (*p*<0.0001, Supplementary Fig 4D), but folic acid still accounted for less than 0.02% of liver folate.

**Figure 5.**
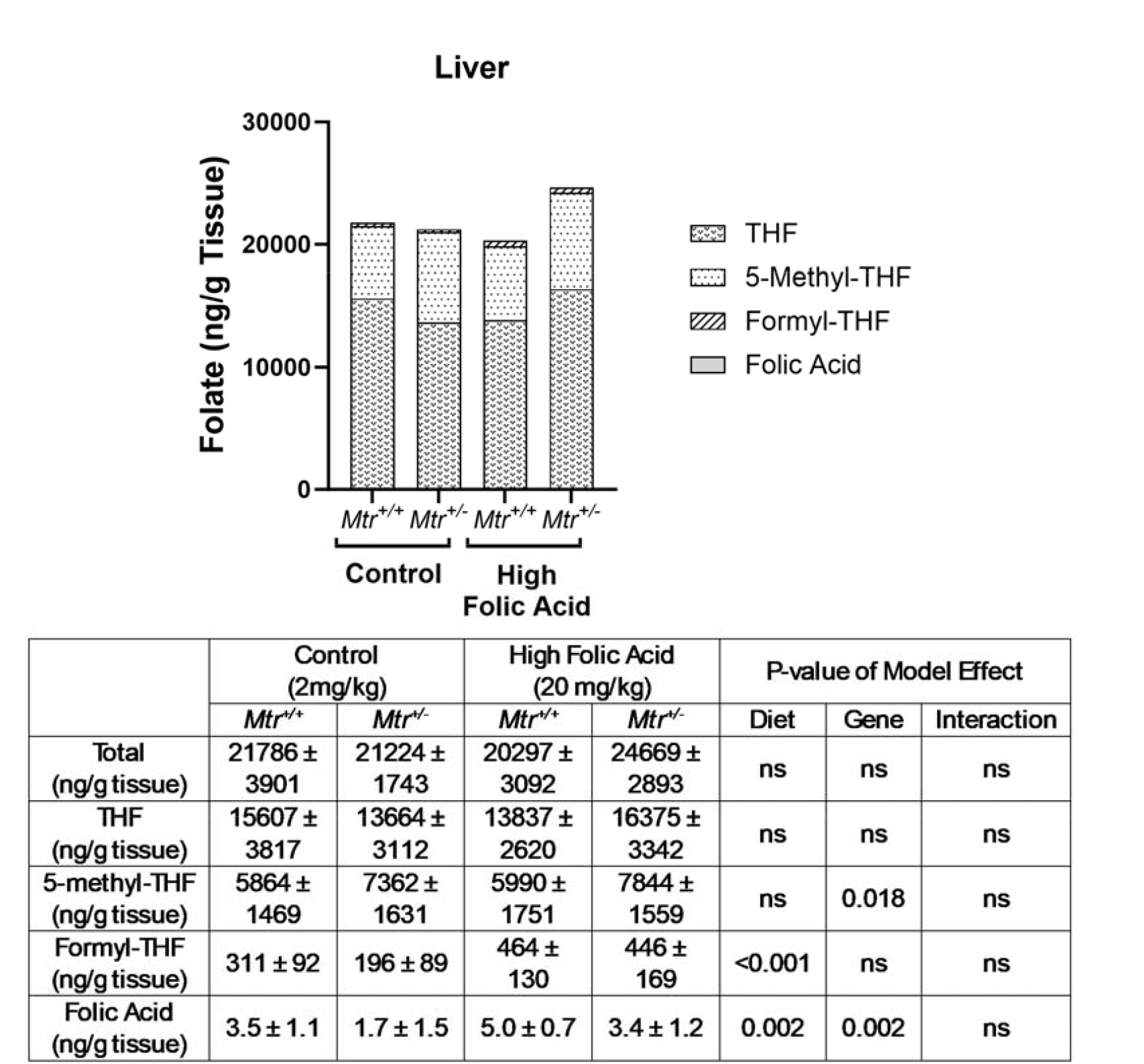
Folate distribution in liver of *Mtr*^*+/+*^ and *Mtr*^*+/-*^ mice fed a control or HFA diet. Folate cofactor distribution in liver was quantified by LCMS/MS. n=6 per group. Two-way ANOVA with Tukey’s post hoc analysis was used to assess main effects of diet and genotype and diet–genotype interaction. Data are presented by group as mean ± SD with statistical significance defined *p* ≤ 0.05.

In *Mtr*^*+/+*^ and *Mtr*^*+/-*^ mice, the main forms of folate present in colon tissue were 5-methyl-THF and formyl-THF (Fig 6). Unlike liver, THF was not detected in colon samples. This agrees with previous findings of cell type-specific distributions of folate forms(21). Folic acid was present in colon tissue, but like liver, it comprised less than 0.2% of total folate present (Fig 6, Supplementary Fig 5C). In agreement with *L. casei* data for colon total folate (Fig 3C), we observed higher total colon folate in mice fed HFA diet when measured by LCMS (*p*<0.0001, Fig 6). Higher total colon folate was driven by higher 5-methyl-THF (*p*<0.0001, Fig. 6) in mice fed the HFA diet. Interestingly, the HFA diet significantly shifted colon folate distribution such that colon 5-methyl-THF was higher (63% vs. 74% of total folate, p<0.0001, Supplementary Fig 5A) and colon formyl-THF was lower (37% vs. 26% of total folate, p<0.0001, Supplementary Fig 5B). A shift in folate distribution and availability of specific folate forms may have implications for FOCM pathway function and cell processes.

**Figure 6.**
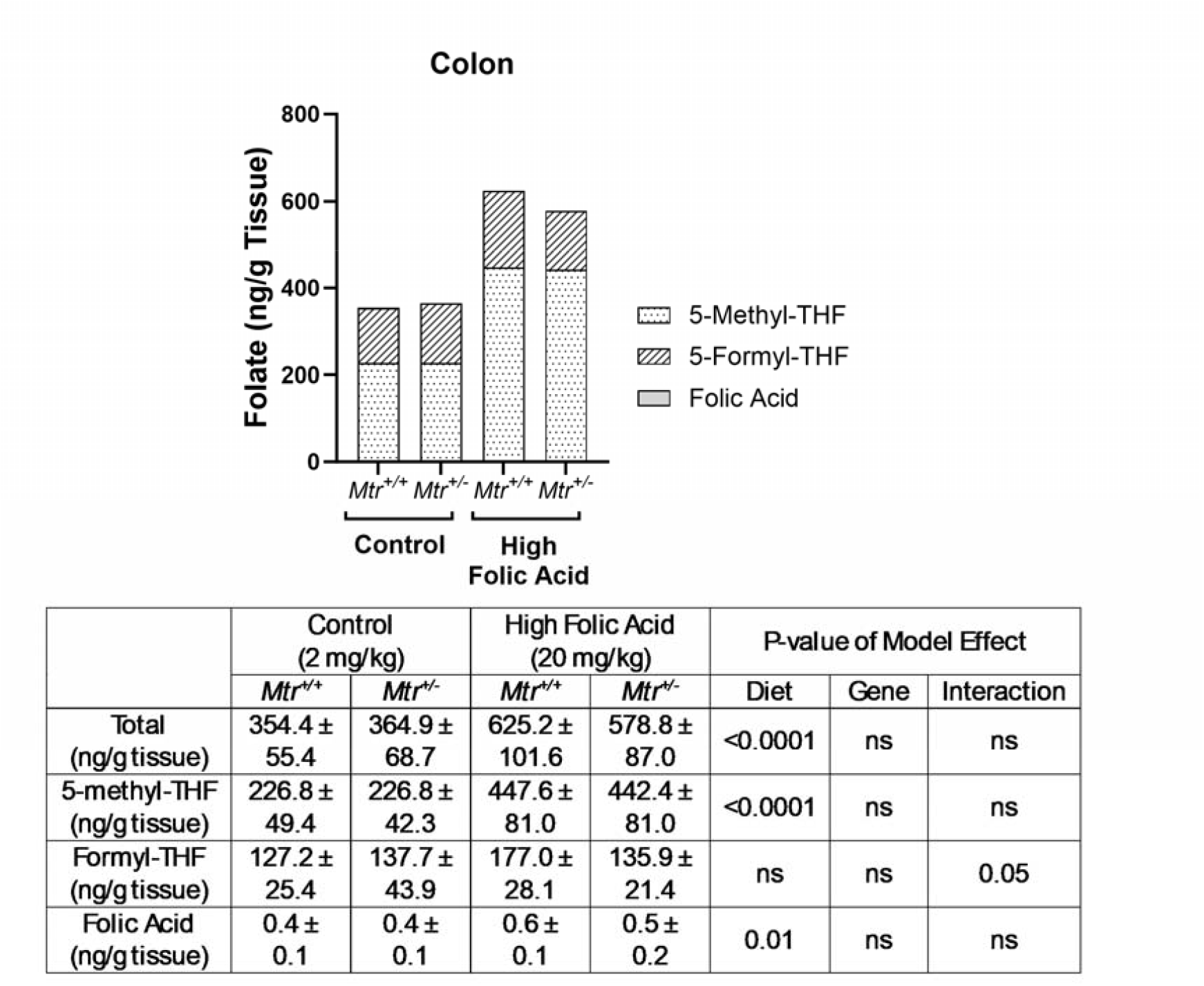
Folate distribution in colon of *Mtr*^*+/+*^ and *Mtr*^*+/-*^ mice fed a control or HFA diet. Folate cofactor distribution in colon was quantified by LCMS/MS. n=6 per group. Two-way ANOVA with Tukey’s post hoc analysis was used to assess main effects of diet and genotype and diet–genotype interaction. Data are presented by group as mean ± SD with statistical significance defined *p* ≤ 0.05.

### High dietary folic acid exposure led to increased uracil misincorporation and γH2AX foci in colon

Impairments in FOCM and folate balance can influence nucleotide synthesis and DNA integrity (22). One marker of disrupted thymidylate synthesis is misincorporation of uracil into genomic DNA (23). Liver genomic uracil content was not affected by exposure to the HFA diet or *Mtr* genotype (Fig 7A). However, both lower *Mtr* and HFA diet resulted in 50% higher colon uracil levels (*p*=0.02 and *p*<0.001, for effects of genotype and diet, respectively, Fig 7D).

**Figure 7.**
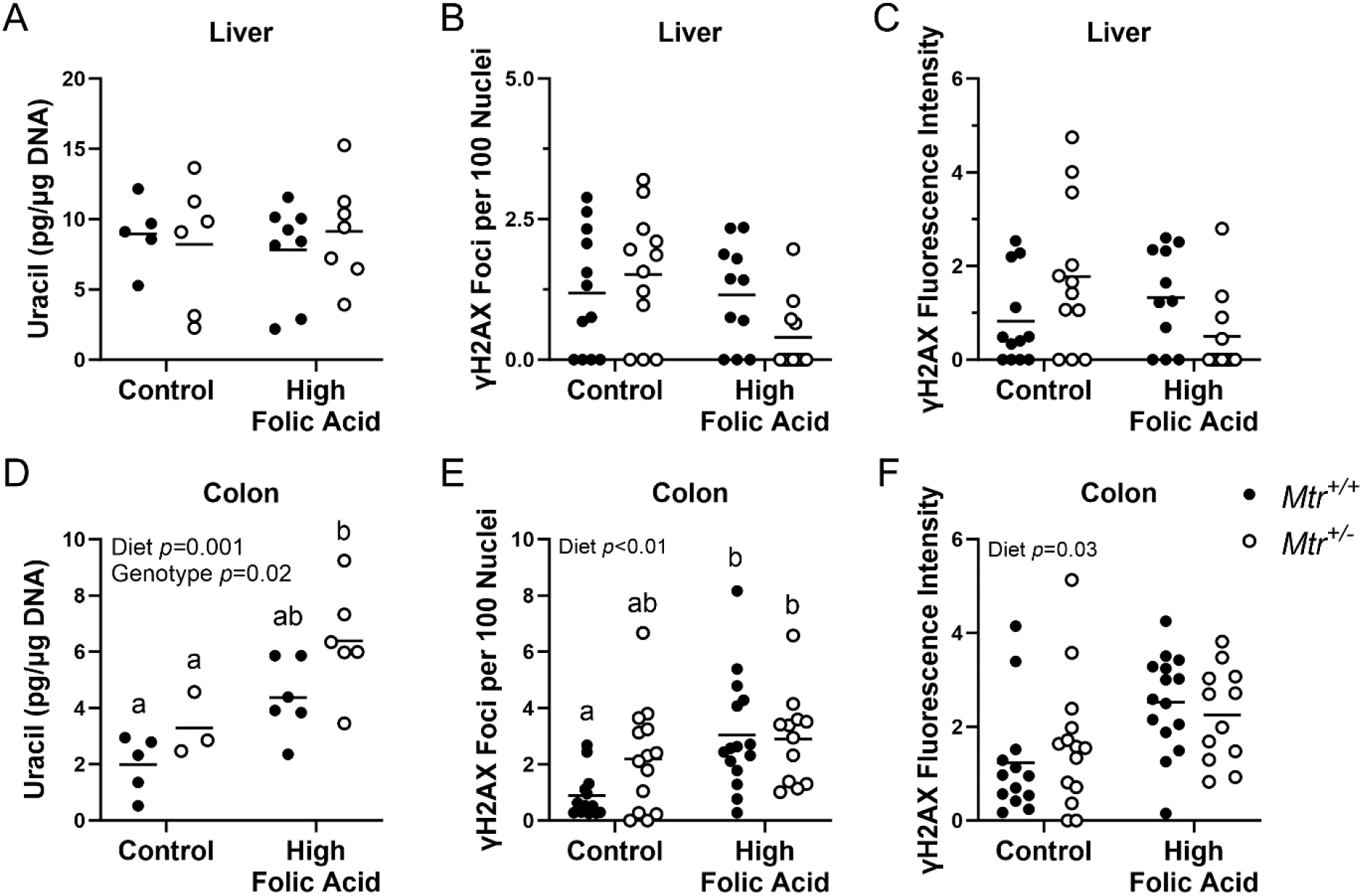
Genomic uracil accumulation and DNA damage (γH2AX) measurements liver and colon from *Mtr*^*+/+*^ and *Mtr*^*+/-*^ mice fed a control or HFA diet. (A) Liver genomic uracil (n=5-8 per group) and (B) colon genomic uracil (n=3-6 per group) measured by GCMS. (C) Liver γH2AX foci (n=3 per group, 4 slides per animal, 3-4 images per slide) and (D) colon γH2AX foci (n=5 per group, 3 slides per animal, 3-4 images per slide) measured by confocal microscopy. (E) Liver γH2AX intensity and (F) colon γH2AX intensity. Uracil datasets were analyzed using a two-way ANOVA with Tukey’s post hoc analysis to assess main effects of diet and genotype and diet–genotype interaction. γH2AX datasets were analyzed using a two-way ANOVA with mixed effects of diet, genotype, and slide. Data are presented as mean with statistical significance defined *p* ≤ 0.05. Groups not connected by a common letter are significantly different.

Uracil misincorporation requires base excision repair enzymes to excise uracil, resulting in an abasic site. Increased rates of uracil misincorporation and subsequent generation of repair-mediated strand breaks may affect genome stability (23). To test if higher uracil content in genomic DNA resulted in higher DNA damage, confocal microscopy was used to measure phosphorylated variant histone H2A (γH2AX) levels, a marker of DNA double-strand breaks and stalled replication forks (21,24). As with genomic uracil levels, liver γH2AX foci count and γH2AX fluorescence intensity were unaffected by exposure to the HFA diet or *Mtr* genotype (Fig 7B-C, Supplementary Fig 6). Notably, there was a modest 2-fold increase in γH2AX foci count and fluorescence intensity in colon with exposure to HFA diet (*p*<0.01 and *p*=0.03, respectively, Fig 7E-F, Supplementary Fig 7). Unlike with colon genomic uracil accumulation, we did not observe a significant effect of *Mtr* genotype on colon γH2AX measurements.

## Discussion

The *Mtr*^*+/-*^ mouse model, which recapitulates the effects of B12 deficiency specific to impaired FOCM in that it increases 5-methyl-THF accumulation (10,19), was used in this study to examine the effects of excess dietary folic acid exposure and to determine if these effects are exacerbated by functional B12 deficiency. Observational studies in humans have found a wide array of negative health outcomes associated with the combination of elevated folate status with low or deficient B12 status. Possible mechanisms for these outcomes have not been investigated in model systems (7). This study is one of the first to investigate this phenomenon in a controlled animal trial. This study demonstrates that 1) high folic acid intake altered folate distribution, nucleotide balance, and DNA damage in a tissue-specific manner, and 2) the combination of high folic acid intake with a functional B12 deficiency did not exacerbate any of these effects.

As expected, excess dietary folic acid exposure elevated both total folate and unmetabolized folic acid in mouse plasma (Fig 2A-C). However, tissue folate content in response to HFA was variable. Liver total folate concentrations were unaffected (Fig 3A), while kidney, colon, and skeletal muscle total folate were higher with HFA diet exposure (Fig 3B-D). This demonstrates the folate content of some tissues is dependent on plasma folate, while others are not. Interestingly, brain total folate concentrations decreased by 10% with exposure to the HFA diet (Fig 3E). The modest reduction in brain folate may not be surprising since both experimental and case studies suggest UMFA in plasma impairs transport of 5-methyl-THF across the blood brain barrier (BBB) (25,26).

DHFR is the enzyme responsible for converting DHF to THF, though it also converts folic acid to DHF, necessary for it to enter the folate cofactor pool. We observed tissue-specific changes in DHFR protein expression in response to the HFA diet. Most notably, DHFR protein levels in colon was 2-3-fold higher with HFA diet (Fig 4D). Brain DHFR protein levels were higher by a modest 30-40% with HFA diet while kidney DHFR levels were unaffected (Supplementary Fig 2A-F). Thus, some tissues may upregulate DHFR protein expression in response to elevated folate and/or folic acid in circulation, but this may be independent of tissue folate accumulation, as not all organs with changes in folate status exhibited changes in DHFR expression.

After observing clear differences in response to the HFA diet depending on tissue type, liver and colon specifically were subjected to further investigation. Liver is a central site of folate storage and one-carbon metabolism homeostasis. Colon epithelia are more mitotically active than liver and may be more sensitive to changes in folate exposure, including impairments in nucleotide synthesis and therefore DNA replication. Moreover, many studies have attempted to investigate the relationship between folic acid exposures and colon cancer incidence in rodent models, though the data are inconsistent (27–30). FOCM produces several one-carbon substituted folate forms that serve as cofactors and a balanced distribution of these folates is essential for supporting nucleotide synthesis. In response to HFA diet, total folate concentration in liver tissue was unaltered while total folate concentration in colon tissue was 2-fold higher compared to mice on C diet. As with total folate concentration in the liver, the distribution of liver folate forms was unaffected by excess dietary folic acid (Fig 5, Supplementary Fig 4). In colon, HFA diet shifted folate distribution to elevate the concentrations of 5-methyl-THF at the expense of formyl-THF (Fig 6, Supplementary Fig 5A-B). Accumulation of 5-methyl-THF can inhibit other FOCM reactions, leaving less folate cofactors available for nucleotide biosynthesis, thus impairing *de novo* dTMP synthesis and subsequent DNA replication and repair(4). Disruption of dTMP synthesis causes accumulation of dUMP and thereafter dUTP(22,23). The increase in dUTP levels facilitates uracil misincorporation (as a “U” base in DNA) because DNA polymerases cannot distinguish between dTTP and dUTP. The shift in colon folate cofactor distribution observed with HFA diet was associated with increased uracil misincorporation in colon genomic DNA (Fig 7D). Uracil misincorporation can be corrected and replaced with thymine via base excision repair. However, an imbalanced ratio of dUTP to dTTP will result in continued misincorporation of uracil and futile cycles of base excision repair, leaving abasic sites that can lead to DNA strand breaks and genomic instability. Previously, impairment of folate-dependent *de novo* dTMP synthesis has been shown to increase levels of both genomic uracil and DNA damage, as measured by phosphorylated histone γH2AX immunostaining (21,24). In the colon, the number of γH2AX foci and γH2AX fluorescence intensity were 2-fold higher in mice fed HFA diet (Fig 7E-F). Since the increase in total folate concentration in colon was driven by an increase in 5-methyHTF, these findings suggest that higher concentration of 5-methyl-THF (induced by exposure to the HFA diet) might impair dTMP synthesis and increase genome instability in colon tissue. Elevated 5-methyl-THF has been shown to suppress *de novo* dTMP synthesis in cultured cells (21). It is important to note that the colon γH2AX levels in mice consuming the HFA diet are orders of magnitude lower than levels seen after exposure to DNA damaging agents (31), but further experiments with genotoxic exposures should be conducted and test whether the effects of such addition exposures they are exacerbated by excess dietary folic acid. However, meta-analyses of human supplementation trials found no evidence that folic acid supplementation was associated with cancer risk (32).

Because both adequate folate and B12 are required for FOCM function, we used the *Mtr*^*+/-*^ mouse model to assess the effects of B12 deficiency specific to FOCM because the *Mtr*^*+/-*^ model results in an increase in 5-methy-THF. The only observed effects of *Mtr* genotype interacting with HFA diet were elevated 5-methyl-THF in plasma (Fig 2B), elevated concentration of 5-methyl-THF in colon tissue (Fig 5), and modestly increased genomic uracil in colon tissue (Fig 7D). Although *Mtr* genotype increased colon uracil content in genomic DNA (Fig 7D), it did not significantly alter colon γH2AX levels (Fig 7E-F, Supplementary Fig 7). *Mtr*^*+/-*^ genotype is a model of mild deficiency that only affects the B12-dependent enzyme of FOCM (19). Therefore, the results presented here suggest that relatively mild, FOCM-specific B12 deficiency does not further exacerbate the effects of excess dietary folic acid on assessed outcomes (folate accumulation, folate distribution, or genome stability). Further investigation with more pronounced B12 deficiency, as would be induced by dietary B12 deficiency, may alter the observed phenotypes. Furthermore, this study focused on exposure to excess dietary folic acid and cannot elucidate the effects of any of high-dose reduced folate intake versus high-dose folic acid intake. This study indicates that the effects of excess dietary folic acid exposure depend on the organ or cell type in question. Investigation of folate distribution, nucleotide synthesis, and DNA damage should be conducted in kidney/brain/other organ systems to further delineate the effects of HFA exposure and if there are any implications for tissue/organ function/disease risk.

## Acknowledgments

The authors acknowledge assistance with statistical analysis from the Cornell Statistical Consulting Unit. The authors also acknowledge the National Defense Science and Engineering Graduate Fellowship program for funding support. The Zeiss LSM 710 Confocal Microscope in the Cornell Institute of Biotechnology’s Imaging Facility was supported by shared instrument grant 1S10RR025502 from the National Institutes of Health.

