## Supplemental materials for "Excess folic acid exposure increases uracil misincorporation into DNA in a tissue-specific manner in a mouse model of reduced methionine synthase expression"

**Supplementary Tables**

| **Dyet #117814GI Custom 2 mg/kg Folic Acid AIN-93G Based Diet** | | | |
| --- | --- | --- | --- |
| **Ingredient** | **kcal/gram** | **grams/kg** | **kcal/kg** |
| Sterile Casein | 3.58 | 200 | 716 |
| L-Cystine | 4 | 3 | 12 |
| Sucrose | 4 | 100 | 400 |
| Cornstarch | 3.6 | 395.5 | 1423.7 |
| Dyetrose | 3.8 | 132 | 501.6 |
| Soybean Oil | 9 | 70 | 630 |
| t-Butylhydroquinone | 0 | 0.014 | 0 |
| Cellulose | 0 | 50 | 0 |
| Mineral Mix #210025 | 0.88 | 35 | 30.8 |
| Folic Acid Premix (1mg/kg Folate) | 4 | 2 | 8 |
| Choline Bitartrate | 0 | 2.5 | 0 |
| Vitamin Mix #317761 (no Folate) | 3.87 | 10 | 38.7 |

**Supplementary Table 1. Composition of control diet (C).** Diets were obtained from Dyets Inc., Bethlehem PA.

| **Dyet #117876GI Custom 20 mg/kg Folic Acid AIN-93G Based Diet** | | | |
| --- | --- | --- | --- |
| **Ingredient** | **kcal/gram** | **grams/kg** | **kcal/kg** |
| Sterile Casein | 3.58 | 200 | 716 |
| L-Cystine | 4 | 3 | 12 |
| Sucrose | 4 | 100 | 400 |
| Cornstarch | 3.6 | 395.5 | 1423.7 |
| Dyetrose | 3.8 | 132 | 501.6 |
| Soybean Oil | 9 | 70 | 630 |
| t-Butylhydroquinone | 0 | 0.014 | 0 |
| Cellulose | 0 | 50 | 0 |
| Mineral Mix #210025 | 0.88 | 35 | 30.8 |
| Folic Acid Premix (1mg/kg Folate) | 40 | 20 | 80 |
| Choline Bitartrate | 0 | 2.5 | 0 |
| Vitamin Mix #317761 (no Folate) | 3.87 | 10 | 38.7 |

**Supplementary Table 2. Composition of high folic acid diet (HFA).** Diets were obtained from Dyets Inc., Bethlehem PA.

**Supplementary Figures**

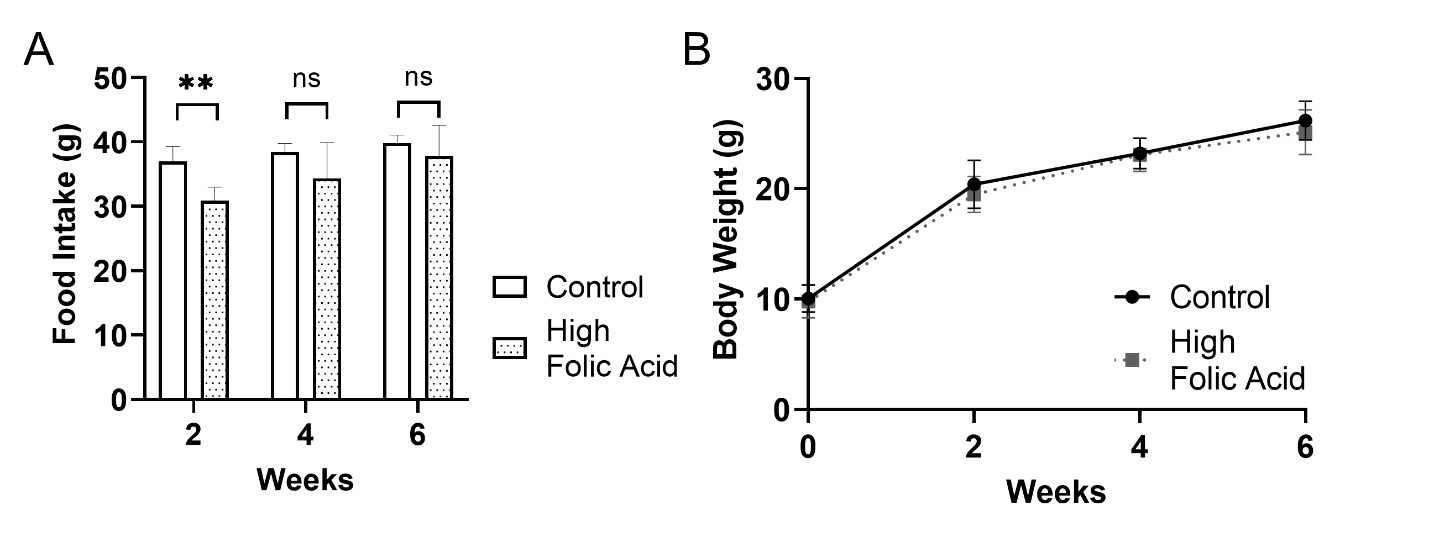

**Supplementary Figure 1. Food Intake and Body Weight of *Mtr* mice fed a control or HFA diet.** (A) Food intake. (B) Body weight. n=10-18 per group. Two-way repeated measures ANOVA with Tukey’s post hoc analysis was used to assess main effects of diet and time and diet–time interaction. Data are presented as mean ± SD with statistical significance defined *p* ≤ 0.05.

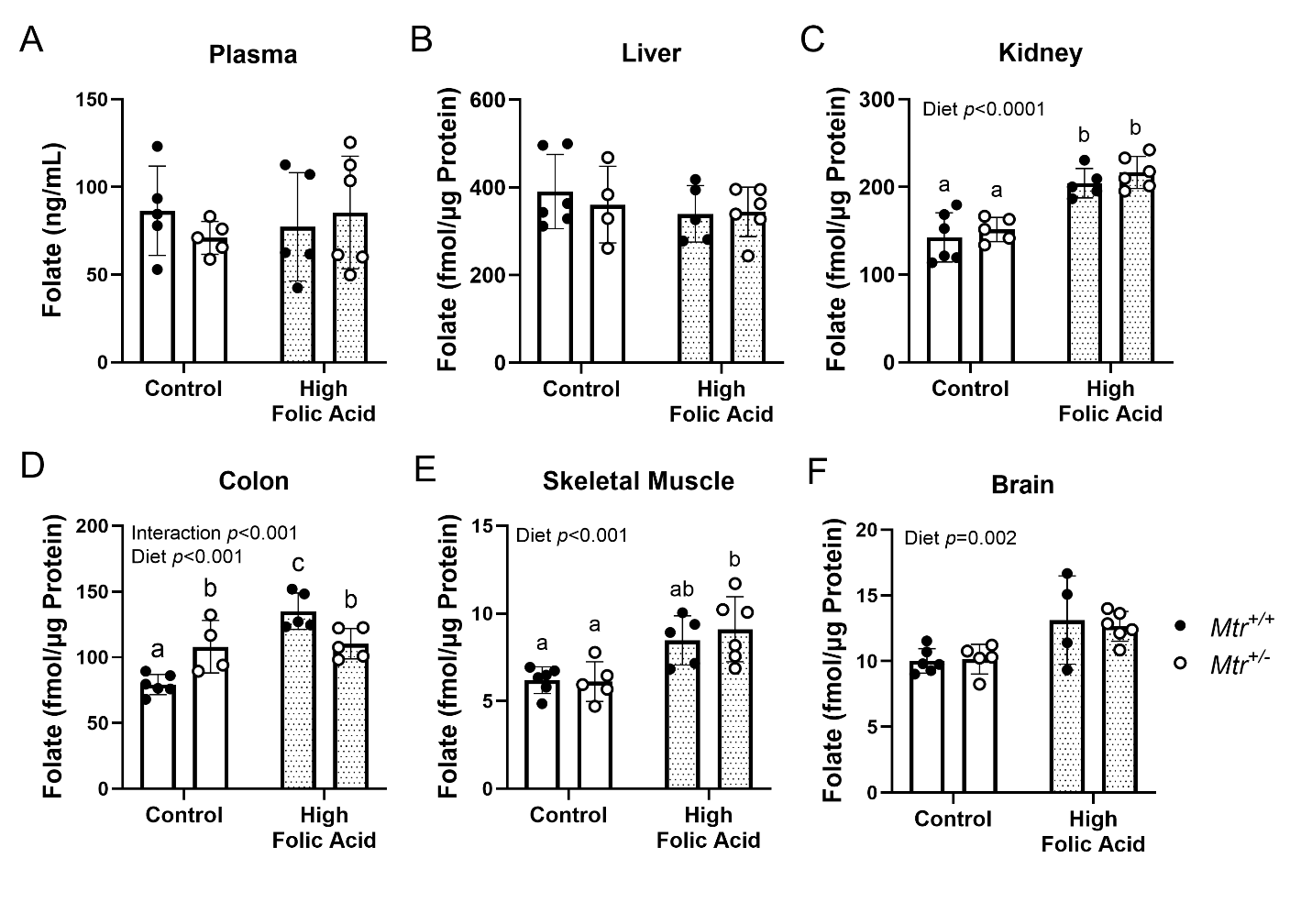

**Supplementary Figure 2. Tissue total folate accumulation in fasted *Mtr^+/+^* and *Mtr^+/-^* mice fed a control or HFA diet.** Measured by L. casei microbiological assay. n=5-6 per group. Two-way ANOVA with Tukey’s post hoc analysis was used to assess main effects of diet and genotype and diet–genotype interaction. Data are presented as mean ± SD with statistical significance defined *p* ≤ 0.05. Groups not connected by a common letter are significantly different.

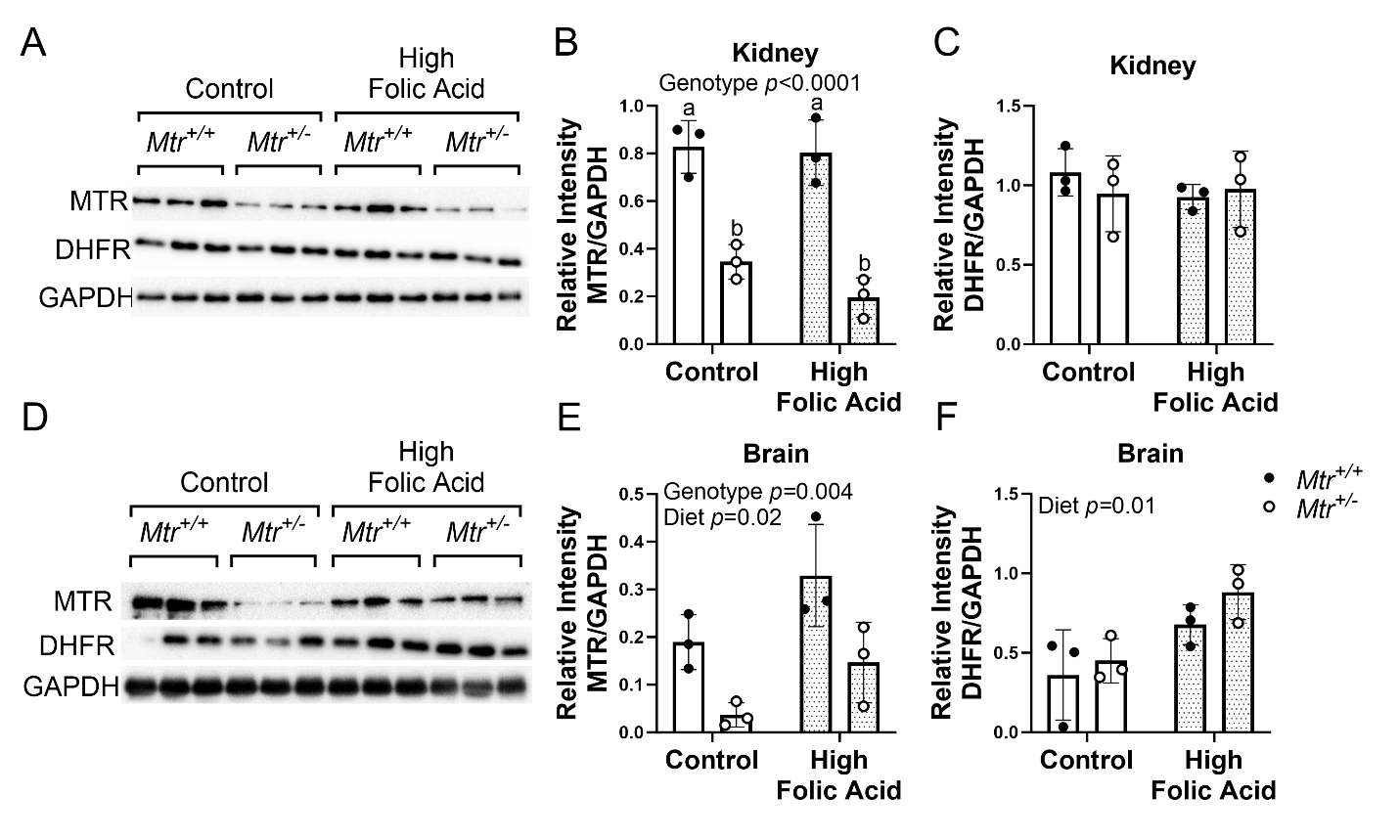

**Supplemental Figure 3. MTR and DHFR protein expression in kidney and brain of *Mtr^+/+^* and *Mtr^+/-^* mice fed a control or HFA diet**. (A) Kidney Blots. (B) Kidney MTR protein levels normalized to GAPDH. (C) Kidney DHFR protein levels normalized to GAPDH. (D) Brain Blots. (E) Brain MTR protein levels normalized to GAPDH. (F) Brain DHFR protein levels normalized to GAPDH. n=3 per group. Two-way ANOVA with Tukey’s post hoc analysis was used to assess main effects of diet and genotype and diet–genotype interaction. Data are presented as mean ± SD with statistical significance defined *p* ≤ 0.05. Groups not connected by a common letter are significantly different. DHFR, dihydrofolate reductase; GAPDH, glyceraldehyde-3-phosphate dehydrogenase; MTR, methionine synthase.

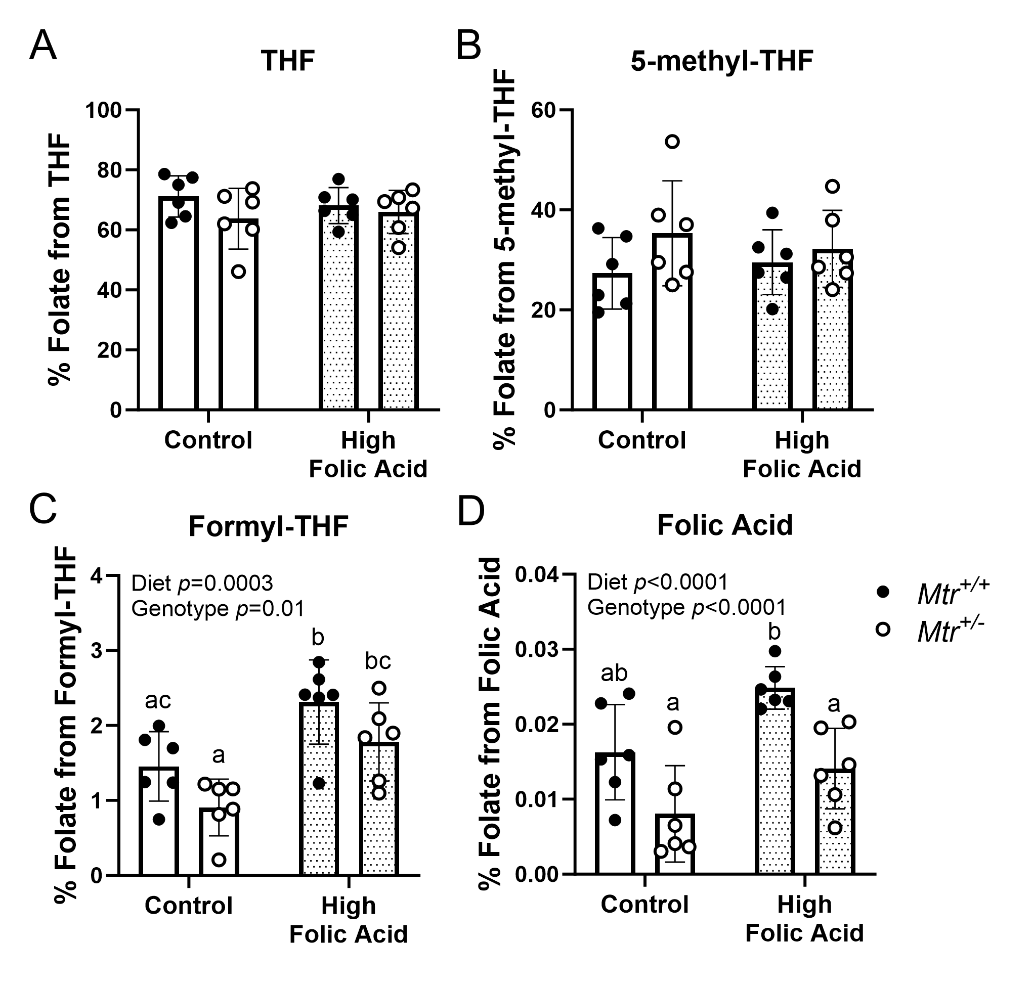

**Supplementary Figure 4. Folate forms relative to total folate in liver of *Mtr^+/+^* and *Mtr^+/-^* mice fed a control or HFA diet**. Folate cofactor distribution was quantified by LCMS/MS. n=6 per group. Two-way ANOVA with Tukey’s post hoc analysis was used to assess main effects of diet and genotype and diet–genotype interaction. Data are presented as mean ± SD with statistical significance defined *p* ≤ 0.05. Groups not connected by a common letter are significantly different. This is the same data as shown in Figure 5 but shown as percentage of total folate.

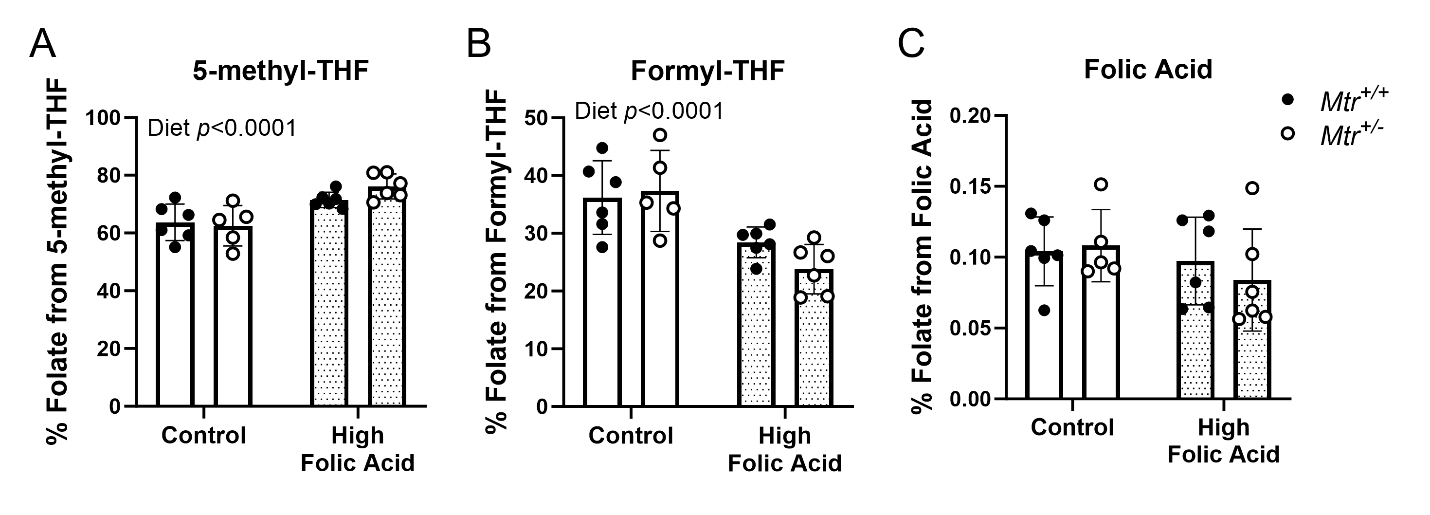

**Supplementary Figure 5. Folate distribution in colon of *Mtr^+/+^* and *Mtr^+/-^* mice fed a control or HFA diet.** Measured by LCMS/MS. n=6 per group. Two-way ANOVA with Tukey’s post hoc analysis was used to assess main effects of diet and genotype and diet–genotype interaction. Data are presented as mean ± SD with statistical significance defined *p* ≤ 0.05. Groups not connected by a common letter are significantly different. This is the same data as shown in Figure 6 but shown as percentage of total folate.

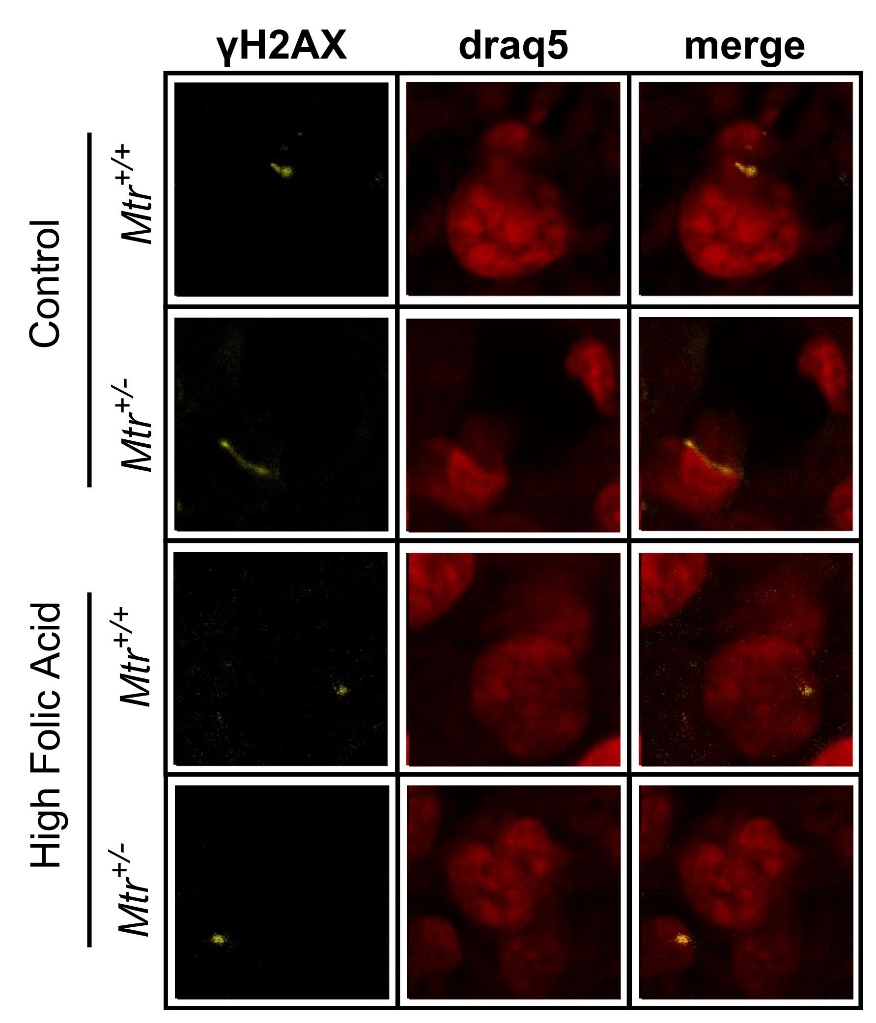

**Supplemental Figure 6. Representative confocal microscopy images used for γH2AX quantification in liver of *Mtr^+/+^* and *Mtr^+/-^* mice fed a control or HFA diet.** draq5 (red, middle) was used as nuclear stain. n=3 animals per group, 4 slides per animal, 3-4 images per slide were acquired and analyzed using ImageJ software.

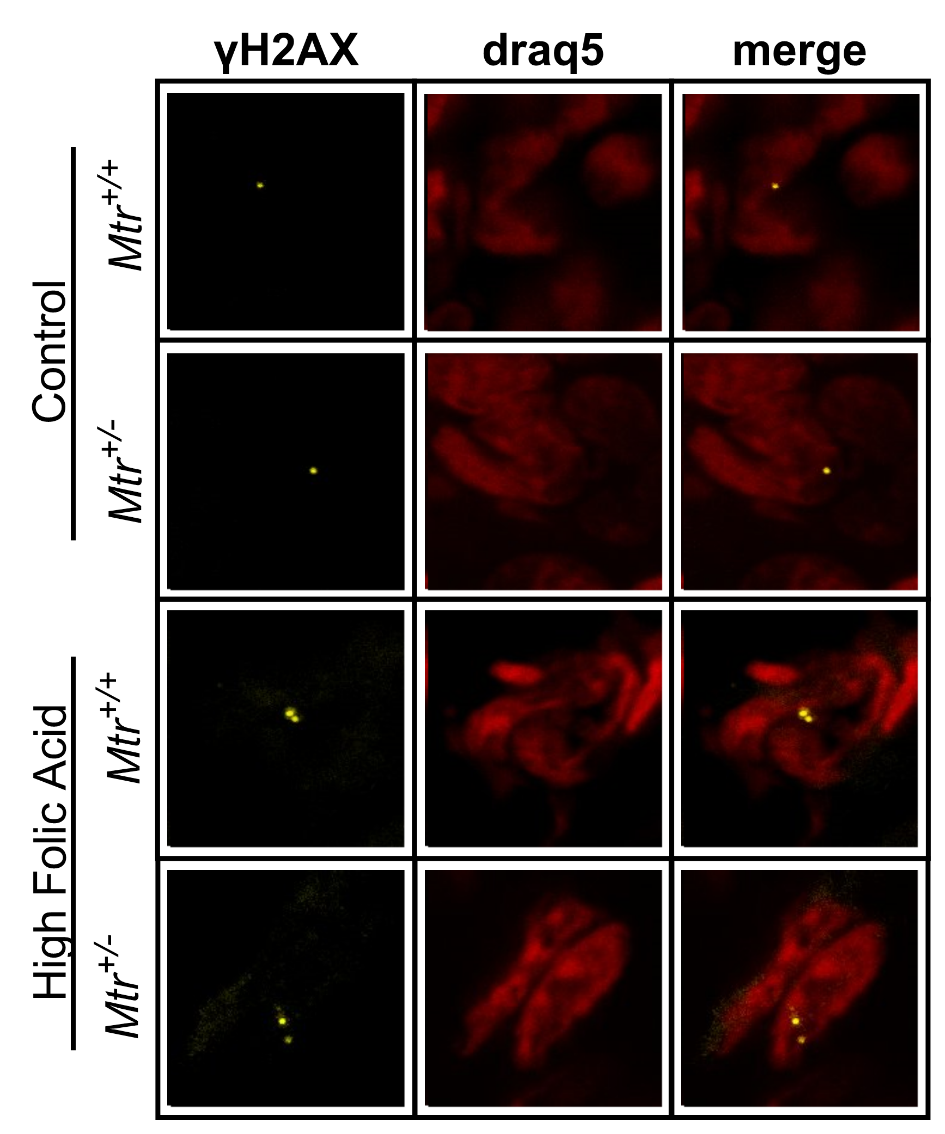

**Supplemental Figure 7. Representative confocal microscopy images used for γH2AX quantification in colon of *Mtr^+/+^* and *Mtr^+/-^* mice fed a control or HFA diet.** draq5 (red, middle) was used as nuclear stain. n=5 animals per group, 3 slides per animal, 3-4 images per slide were acquired and analyzed using ImageJ software.
